# A Vascular Dissection and Rupture Linked Metabolite Acts Via BLT2 Receptor

**DOI:** 10.1101/2024.02.15.580482

**Authors:** Yuyu Li, Jiaqi Yu, Weiyao Chen, Xin Tan, Xuan Xu, Rui Lin, Xue Wang, Wenxi Jiang, Jie Du, Yuan Wang

**Affiliations:** Beijing Anzhen Hospital, Capital Medical University; Key Laboratory of Remodeling-related Cardiovascular Diseases, Ministry of Education; Beijing Collaborative Innovation Centre for Cardiovascular Disorders, No. 2 Anzhen Road, Chaoyang District, Beijing 100029, China; Beijing Institute of Heart, Lung and Blood Vessel Disease, No. 2 Anzhen Road, Chaoyang District, Beijing 100029, China

**Author notes:** These authors contributed equally to this work (Yuyu Li, Jiaqi Yu, Weiyao Chen). **Correspondence** Correspondence to: Yuan Wang, Beijing Anzhen Hospital, No. 2 Anzhen Rd, Chaoyang District, Beijing 100029, China; or Jie Du, PhD, Beijing Anzhen Hospital, No. 2 Anzhen Rd, Chaoyang District, Beijing 100029, China. or.

**Keywords:** aortic dissecting, macrophage, extracellular matrix, muscle, smooth, vascular

## Abstract

**BACKGROUND:** Thoracic aortic dissection (TAD) is a life-threatening vascular disease that requires effective drug treatment to prevent progression and rupture. Because arachidonic acid metabolism is involved in inflammation and vascular homeostasis, we investigated the roles of arachidonic acid metabolites in TAD pathogenesis and their utility as therapeutic targets.

**METHODS:** Serum metabolomics analysis was performed to characterize arachidonic acid metabolites in TAD patients and a TAD mouse model. 12/15-LOX expression was profiled in the aortic tissues of TAD patients and the TAD mouse model. Four-week-old male Alox15 knockout mice (Alox15^−/−^), 12-HETE-treated mice, ML351 (12/15-LOX inhibitor)-treated mice, and LY255283 (leukotriene B 4 receptor 2 [BLT2] antagonist)-treated mice received β-aminopropionitrile monofumarate (BAPN, 1 g/kg/day) for 4 weeks to model TAD, then underwent assessment of TAD progression. Interaction of 12-HETE produced by macrophages with BLT2 receptor-expressing cells was detected by molecular docking and immunoblotting.

**RESULTS:** Serum levels of 12-HETE and the expression of 12/15-LOX in aortic tissue were significantly increased in TAD patients and BAPN-treated TAD mice. BAPN-induced TAD progression was significantly ameliorated in Alox15-deficient or -suppressed mice. 12-HETE directly interacted with BLT2 receptors on macrophages, activating the downstream NOX-1/ROS/NF-κB signaling pathway to induce inflammatory cytokine release. This initiated inflammatory cell recruitment and exacerbated extracellular matrix degradation, leading to phenotype switching in vascular smooth muscle cells (VSMCs). Additionally, treatment with ML351 and LY255283 significantly reduced the rates of dissection rupture and combined treatment could maximize the curative effect.

**CONCLUSIONS:** 12-HETE may amplify the inflammatory cascade and trigger aberrant phenotype switching in VSMCs during TAD development. The reduction of circulating 12-HETE or antagonism of its receptor may be new targets for TAD prevention and treatment.

**Clinical Perspective:** *What Is New?:*

- The expression levels of 12/15-LOX and its metabolite 12-HETE were elevated in TAD patients and TAD mice.
- Increased levels of 12-HETE directly bind to BLT2 receptors in macrophages, thereby initiating inflammatory cascades that downregulate VSMC differentiation markers through the suppression of IL-6.
- Deletion or pharmacologic inhibition of 12/15-LOX and suppression of BLT2 mitigated TAD development by alleviating inflammation and VSMC phenotype switching.

*What Are the Clinical Implications?:*

- The inhibition of 12-HETE-related pathways, through mechanisms such as reducing the plasma 12-HETE content or blocking its receptor, may represent a novel therapeutic strategy for TAD.
- Further studies are needed to explore the diagnostic value of serum 12-HETE as a novel biomarker for TAD.

## Introduction

Thoracic aortic dissection (TAD) is a severe medical condition with rapid progression and high morbidity in the acute phase.^1, 2^ Although surgical therapies have significantly improved in recent years, there are insufficient clinically effective medications to slow or prevent aortic degeneration.^3^ Therefore, investigations of TAD pathogenesis are needed to develop effective treatment strategies.

TAD pathogenesis is complex-it involves inflammation, apoptosis, smooth muscle cell phenotype switching, and extracellular matrix degradation.^4, 5^ Vascular smooth muscle cells (VSMCs) comprise the majority of cellular components in the middle aorta; their phenotypes range from a quiescent (contractile) state (responsible for regulating blood flow and pressure) to a proliferative (synthetic) phenotype (capable of synthesizing matrix metalloproteinase (MMP)-2 and MMP9).^6^ Aberrant VSMC phenotype switching is triggered by environmental stimuli, such as inflammatory cytokines and reactive oxygen species (ROS).^7^ In response to inflammatory cues, VSMCs undergo dedifferentiation to a synthetic phenotype, which exhibits enhanced proliferative and migratory capacities.

Arachidonic acid metabolism is a potent regulator of inflammation.^8^ Arachidonic acid, typically distributed in the cell membrane in phospholipid form, is released in response to pathological stimuli. The released arachidonic acid is converted into eicosanoids (prostaglandins and leukotrienes), thromboxanes, and other signaling lipid metabolites that may trigger inflammatory responses.^9, 10^ Bioactive metabolites of arachidonic acid can also trigger cardiomyocyte ferroptosis, thereby promoting myocardial ischemia-reperfusion injury.^11, 12^ There is experimental evidence that arachidonic acid and its metabolites are involved in pathological processes including oxidative stress, cardiomyocyte apoptosis, and platelet hyperactivation.^13, 14^ Thus, they play important roles in cardiovascular diseases such as myocardial infarction and hypertension. However, the potential contributions of these eicosanoids to TAD development have not been investigated. We speculated that the coordinated release of arachidonic acid metabolites after vascular injury initiates an inflammatory cascade and subsequent VSMC phenotype switching, which may be an important contributing factor in TAD formation.

Here, we performed an arachidonic acid-targeted metabolomics analysis using serum samples collected from TAD patients and healthy controls. We observed substantial accumulation of LOX family-associated metabolites in TAD serum, including 12-hydroxyeicosatetraenoic acid (12-HETE), a product of arachidonate 12/15-lipoxygenase (12/15-LOX). We found that 12/15-LOX was increased in diseased aortic tissue from TAD patients and TAD mice. Our observations in an animal model validated this highly conserved role for 12/15-LOX in TAD. Furthermore, we found that 12-HETE, detected in early vascular injury, induces aortic inflammation by activating BLT2 and promoting aberrant VSMC phenotype switching. Overall, we identified 12-HETE as a key initiator of the inflammatory response that promotes VSMC phenotype switching during TAD formation, then demonstrated its utility as a therapeutic target for TAD prevention.

## METHODS

The data that support the findings of this study are available from the corresponding author on reasonable request. Expanded Methods are provided in the Supplemental Material.

### Study population, Human Tissue Collection and Ethics Statement

Participants were enrolled in the DPANDA registry study for aortic aneurysm and/or dissection. Experienced cardiothoracic surgeons confirmed the presence of TAD during surgery, and the clinical phenotype diagnosis was verified by standard histopathology at the Beijing Anzhen Hospital Affiliated to Capital Medical University. Age-matched healthy individuals were included for serum collection. Characteristics of patients, controls, and healthy individuals are presented in Supplemental Table I. Dissecting aortic aneurysmal tissue and sera were collected during surgery from patients undergoing aortic root and ascending aorta replacement. Control aortic tissues were obtained from age-matched patients undergoing heart transplant surgery without aortic aneurysm, dissection, coarctation, or previous aortic repair. The study received approval from the Ethics Committee of the Beijing Anzhen Hospital Affiliated to Capital Medical University (ClinicalTrials.gov: NCT03233087) and was conducted in accordance with the Declaration of Helsinki. Written informed consent was obtained from all patients.

### Animals

The animal experiments were carried out in specific pathogen-free barrier conditions, adhering to institutional guidelines. Approval for the protocol was obtained from the Committee on the Ethics of Animal Experiments of Capital Medical University. Further details regarding the animals used can be found in the Supplementary Material (Figure S1).

## RESULTS

### 12/15-LOX Expression and Serum 12-HETE Levels Were Elevated in TAD Patients and TAD Mice

To identify metabolites potentially associated with TAD, we performed arachidonic acid-targeted metabolomics analyses of 47 plasma samples from TAD patients and 39 plasma samples from healthy controls. Furthermore, we analyzed 27 plasma samples from β-aminopropionitrile monofumarate (BAPN)-treated TAD mice and 16 plasma samples from sham mice (Figure S 2A). In total, 74 metabolites were detected including 4 polyunsaturated fatty acids (ARA, EPA, DPA, DHA) and 70 eicosanoids across the cyclooxygenase (COX), lipoxygenase (LOX) and cytochrome P450 (CYP450) pathways. Furthermore, partial least square discriminant analysis, a mathematical procedure that decreases data dimensionality while preserving most of the variance by orthogonal transformation,^15^ indicated that AA metabolite variables were distinguishable between TAD patients and healthy controls (Figure 1A) and between BAPN-treated TAD and sham mice (Figure 1F). Among these AA metabolites, 12-HETE, implicated as a primary metabolite in the 12/15-LOX pathway,^16^ exhibited the greatest alteration and was identified as the main contributor to the overall difference in serum samples from human participants (Figure 1B, Figure S2B) and mice (Figure 1G, Figure S2E). Additionally, relative quantification of 12-HETE by mass spectrometry showed that serum concentrations were significantly greater in TAD patients than in healthy controls (Figure 1C). This finding was validated in BAPN-treated TAD mice (Figure 1H). Besides, serum 12-HETE concentrations allowed differentiation of patients with TAD from healthy controls, with an AUC of 0.995(0.987-1.000) (Figure S 2C) and higher level of 12-HETE was observed in TAD patients experienced major adverse events (Figure S2D). Heatmap analysis of the levels of 74 metabolites revealed considerable differences between TAD patients and healthy controls (Figure 1D and 1I). Western blotting (WB) and immunofluorescence (IF) staining analysis showed that 12/15-LOX expression in aortic tissues was significantly greater among TAD patients than among healthy controls (Figure 1K and 1M). Increased expression of 12/15-LOX was also observed in BAPN-treated TAD mice (Figure 1L and 1N, Figure S2F-S2G). WB quantification of 12/15-LOX is depicted in Figure 1E and 1J. These results suggest that 12/15-LOX and its metabolite 12-HETE have roles in TAD development.

**Figure 1.**
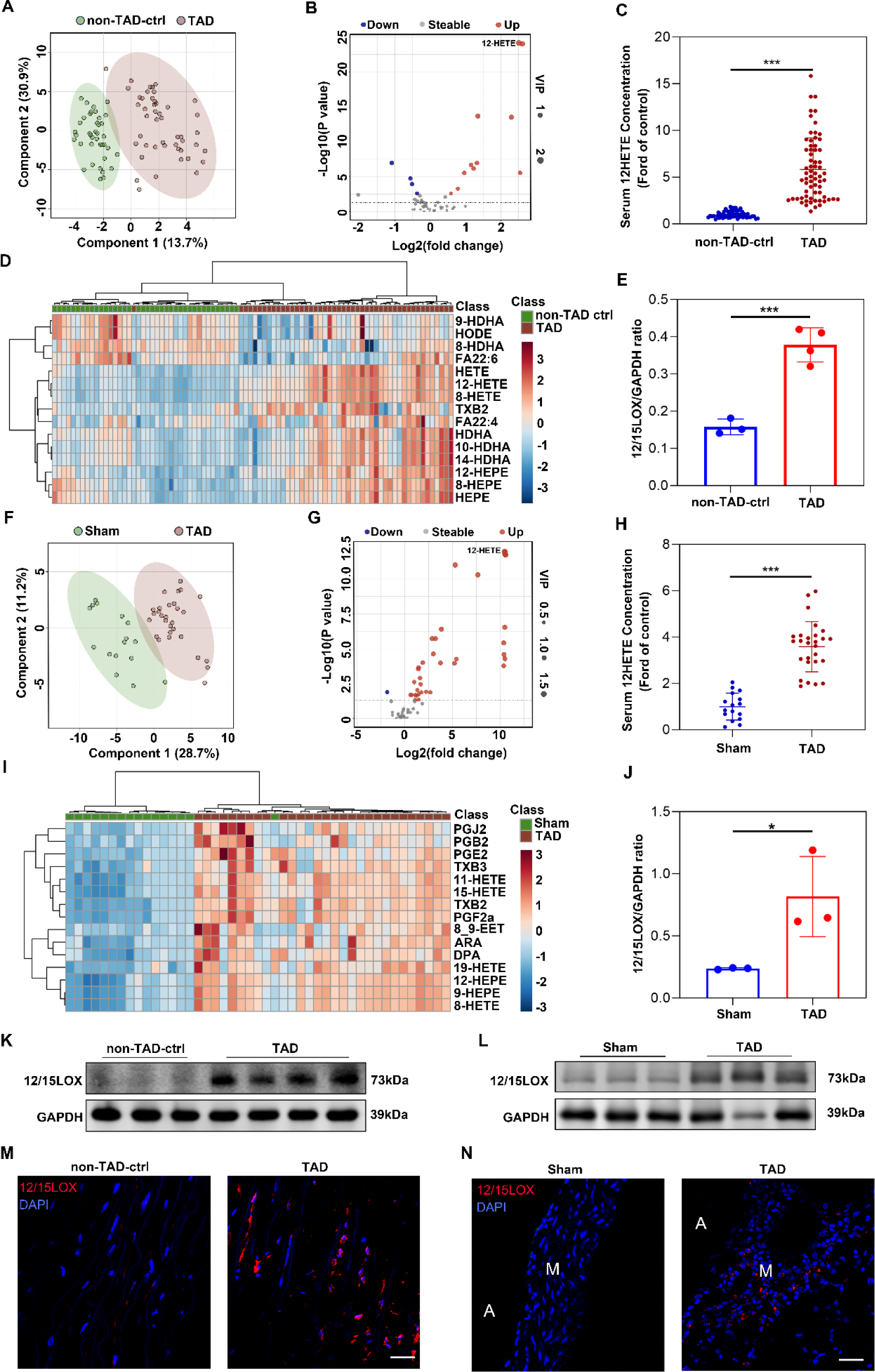
Metabolomics profiling identifies ALOX15-15-HETE as the most outstanding axis in TAD. **A**, AA metabolites were extracted and interpreted by principal component analysis (PCA), and the 2D score plots display repertoires of non-TAD control and TAD patients. Each point represents a sample, and ellipses represent 95% confidence regions (n=39 and 47). **B,** Volcano plot shows − log10(P-value) on the y-axisversuslog2(fold change) on the x-axis. Each point represents a different metabolite and the greater the scattered point, the greater the value of variable importance is in the projection (VIP). **C,** Serum 12HETE concentration (ford of control) of TAD individuals and non-TAD controls. **D,** Serum contents of AA pathway metabolites of TAD individuals and non-TAD controls based on targeted metabolomics. **E,** Western blot analysis and quantification of 12/15LOX expressed in aorta of controls without TAD and patients with TAD (n=3 and 4 per group). **F,** AA metabolites were extracted and interpreted by PCA, and the 2D score plots display repertoires of healthy control and TAD mice. Each point represents a sample, and ellipses represent 95% confidence regions (n=16 and 27) **G,** Volcano plot of AA metabolites from the serum of healthy control and TAD mice. **H,** Serum 12HETE concentration of healthy control and TAD mice. **I,** Serum contents of AA pathway metabolites of healthy control and TAD mice based on targeted metabolomics. **K,** Western blot analysis of 12/15LOX levels in aorta of controls without TAD and patients with TAD (n=3 and 4 per group). **L**, Western blot analysis of 12/15LOX levels in healthy control and TAD mice (n=3 per group).**M**, Representative images of human aorta stained with 12/15LOX (12/15LOX; red) and DAPI (blue). Scale bars = 50μm; media (M). **N,** Representative images of mouse aorta stained with 12/15LOX (12/15LOX; red) and DAPI (blue). Scale bars = 50μm; media (M); adventitia. \**p* < 0.05, \*\**p* < 0.01 and \*\*\**p* < 0.001, n.s. = not significant, one-way ANOVA, Tukey’s multiple comparisons test.

### Global Alox15 Knockout Mitigates TAD Development in Mice

To investigate the role of 12-HETE in aortic dissection, we performed in vivo experiments using a mouse model of BAPN-induced TAD. First, global 12/15-LOX knockout mice (Alox15^−/−^) were purchased from the Jackson Laboratory. Next, 4-week-old male 12/15-LOX knockout mice (n=20) and C57BL/6J wild-type (WT) mice (n=22) were treated with BAPN for 28 days. During the 28 days of BAPN administration, 68% (n=15) of the C57BL/6J WT mice and 30% (n=7) of the Alox15^−/−^ mice died of aortic dissection and rupture (Figure 2B and 2C). 27% of the surviving WT mice and 20% of the surviving Alox15^−/−^ mice developed aortic dissection after BAPN treatment (Figure 2A and 2C). Vascular ultrasound images and measurements of maximum aortic diameter on day 28 after modeling also demonstrated that Alox15 knockout mitigated BAPN-induced aortic dilation compared with WT controls (Figure 2D and 2E). Hematoxylin and eosin (HE) staining and Elastica van Gieson (EVG) staining showed that dissecting aneurysm formation and elastin disruption were also alleviated in BAPN-treated Alox15^−/−^ mice compared with C57BL/6J mice (Figure 2F). In the absence of BAPN treatment, WT and Alox15^−/−^ mice exhibited minimal differences in aortic wall thickness, diameter, and TAD formation (Figure S3A–S3C). Body weight, systolic pressure, and diastolic pressure showed minimal differences between Alox15^−/−^ mice and WT controls (Figure S3D–S3F). These findings suggested that 12/15-LOX deficiency can reduce TAD formation.

**Figure 2.**
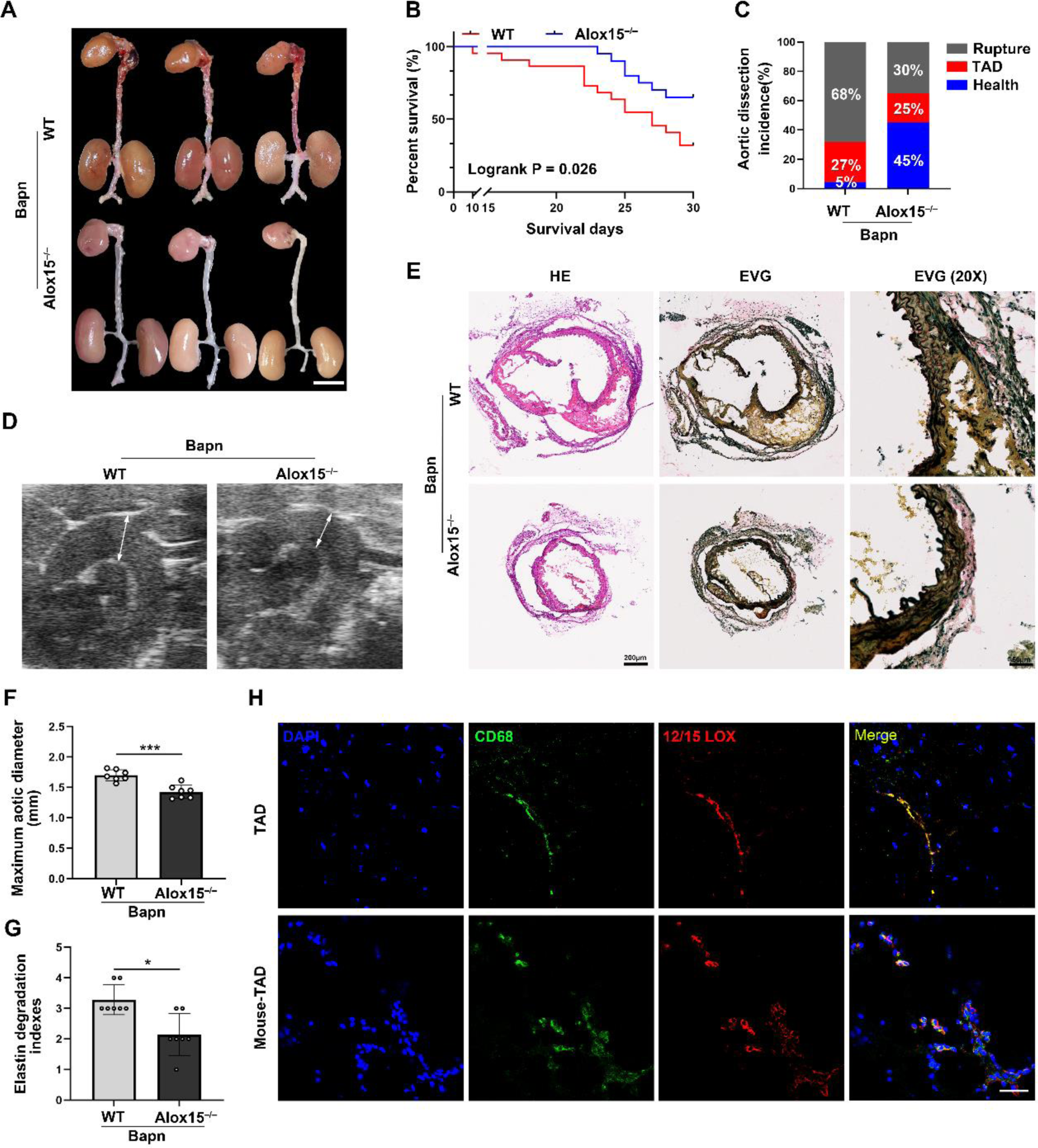
Global Alox15 knockout mitigates TAD development in mice. **A - F**, Wild-type (WT) and Alox15^−/−^ mice were treated with BAPN (β-aminopropionitrile monofumarate) for 28 days. **A**, Representative macrographs of aorta (scale bar = 5 mm). **B**, Survival rate was estimated by Kaplan-Meier method and compared by log-rank test (n=22, 20). **C**, Thoracic aortic dissection (TAD) incidence. **D**, Representative ultrasound images of thoracic aorta. **E**, Representative macroscopic images of aorta sections stained with hematoxylin and eosin (HE) and elastic–Van Gieson (EVG) (scale bars, 200 μm; 50 μm). **F**, Measurements of maximum aortic diameter (n=6 per group). **G,** Elastin break grades were analyzed by Kruskal-Wallis followed by Dunn multiple comparisons test. **H**, Representative confocal images of 12/15LOX (red) colocalized with CD68 (green)–positive macrophages in aorta tissues from patients with thoracic aortic dissection (TAD) and TAD mice. (scale bars, 50 μm). \**p* < 0.05, \*\**p* < 0.01 and \*\*\**p* < 0.001, n.s. = not significant, one-way ANOVA, Tukey’s multiple comparisons test.

### Alox15 Deficiency Abolishes Injury-Induced Contractile-to-Synthetic Phenotype Switch in VSMCs

Considering that the contractile-to-synthetic phenotype switch in VSMCs is a key pathogenic component of TAD development, we measured the expression levels of aorta-associated proteins such as α-smooth muscle actin (α-SMA), smooth muscle 22α (SM22α), and collagen I. WB analysis revealed that Alox15 deletion had minimal effects on basal levels of contractile markers such as SM22α and α-SMA. After BAPN treatment, WT mice exhibited a significant decrease in contractile markers in the aorta, whereas Alox15 deletion rescued these levels (Figure 3A and 3C–3F). IF staining demonstrated that Alox15 deletion counteracted undesirable BAPN-induced changes in SM22α and α-SMA (Figure 3G–3I). Moreover, the protein levels of MMP2 and MMP9, which promote TAD formation, were considerably higher in TAD tissues than in adjacent non-TAD tissues. Additionally, MMP2 and MMP9 were upregulated in the aorta of Alox15^−/−^ TAD mice, indicating that extracellular matrix formation was inhibited in such mice (Figure S4, Figure 3J and 3K). Those results suggested that Alox15 deficiency can activate differentiation signaling in VSMCs and inhibit extracellular matrix degradation.

**Figure 3.**
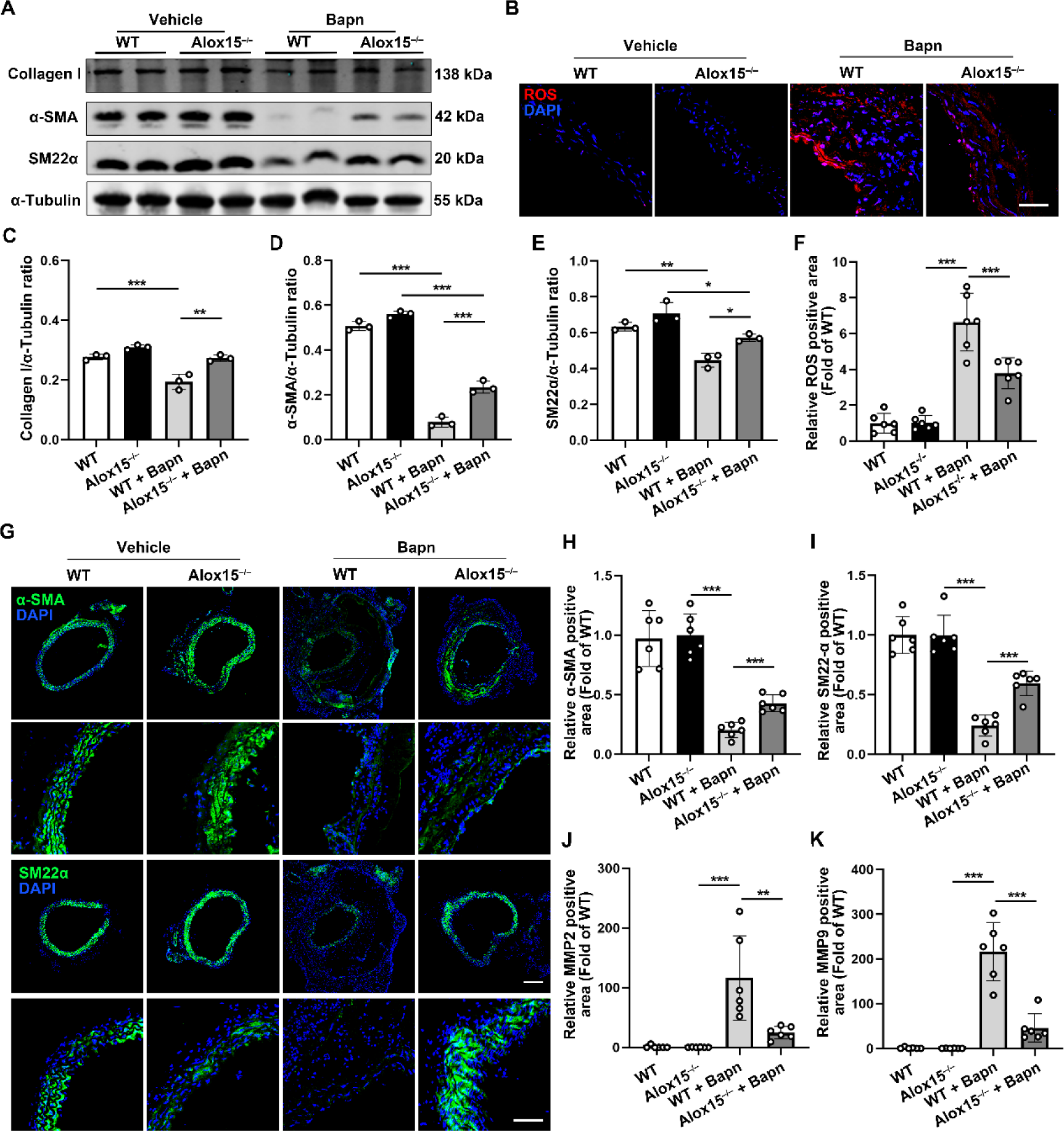
Alox15 deficiency abolishes injury-induced contractile-to-synthetic phenotype switch in VSMCs. **A**–**K,** Wild-type (WT) and Alox15^−/−^ mice were treated with or without BAPN for 28 days. **A**, Representative Western blot images of Collagen I, α-SMA and SM22α in VSMCs. **B**, The level of reactive oxygen species (ROS) in the aorta section of BAPN - and saline-treated WT and Alox15^−/−^ mice were evaluated by dihydroethidium (DHE) staining and quantified by determining the ratio of DHE-positive area (n = 6) (scale bars, 50μm) (magnified photographs). **C–F**, Western blot analysis and quantification of Collagen I, α-SMA and SM22α expression in the aorta of BAPN - and saline-treated WT and Alox15^−/−^ mice (n=6 per group; unpaired 2-tailed t test for SM22α and α-SMA. **G**, Immunofluorescence staining of SM22α (Green) and α-SMA (Green) in aorta. Nuclei were counterstained with DAPI (blue; scale bars, 200 μm, 50 μm). **H** and **I**, Quantification of SM22α and α-SMA– positive area in aorta. **J** and **K,** Quantification of MMP2 and MMP9– positive area in aorta. \**p* < 0.05, \*\**p* < 0.01 and \*\*\**p* < 0.001, n.s. = not significant, one-way ANOVA, Tukey’s multiple comparisons test.

Considering that ROS are key signaling molecules with important roles in the progression of inflammatory disorders, we examined ROS levels in the aortas of BAPN- and saline-treated mice. As shown in Figure 3B, Alox15 deletion alleviated ROS production in the aorta of BAPN-treated TAD mice. Taken together, these results suggest that Alox15 silencing is associated with decreased formation of BAPN-induced TAD.

### Alox15 Deficiency Reduces BAPN-Induced Vascular Inflammation in Mice

Considering that Alox15 deletion can alleviate ROS production in the aorta of TAD mice, we evaluated its effect on the inflammatory response in BAPN-treated TAD mice. IF staining revealed fewer Ly6G^+^ neutrophils and CD68^+^ macrophages infiltrating the aorta in Alox15^−/−^ TAD mice than in WT TAD mice (Figure 4A and 4B). Furthermore, the expression levels of inflammatory factors interleukin (IL)-1β, IL-6, tumor necrosis factor (TNF)-α, and CCL2 were decreased in serum samples from Alox15^−/−^ mice (Figure 4C). There were positive correlations between plasma 12-HETE and the above inflammatory factors (Figure 4G). It’s worth noting that elevated 12-HETE was observed in mice treated with BAPN for 3w, precedes the increase in IL-6, IL-1β and CCL2 (n=5,8 and 8, Figure S5). Moreover, flow cytometry analysis indicated that there were considerably fewer neutrophils and macrophages in Alox15^−/−^ mice (Figure 4D–4F). What’s more, Alox15 deletion in myeloid cells resulted in dramatic suppression of myelopoiesis in the bone marrow (BM). There were fewer newly formed neutrophils and macrophages in the BM (Figure S6), spleen (Figure S7), and blood (Figure 4E and 4F). Further, we also performed correlation analyses between serum 12HETE levels and serum inflammatory factor (IL-1β, IL-6, TNF-α and CCL2) levels, and the results showed that 12-HETE was positively correlated with these pro-inflammatory factors (Figure 4G). These findings demonstrated that 12/15-LOX deficiency had a critical role in regulating TAD inflammation.

**Figure 4.**
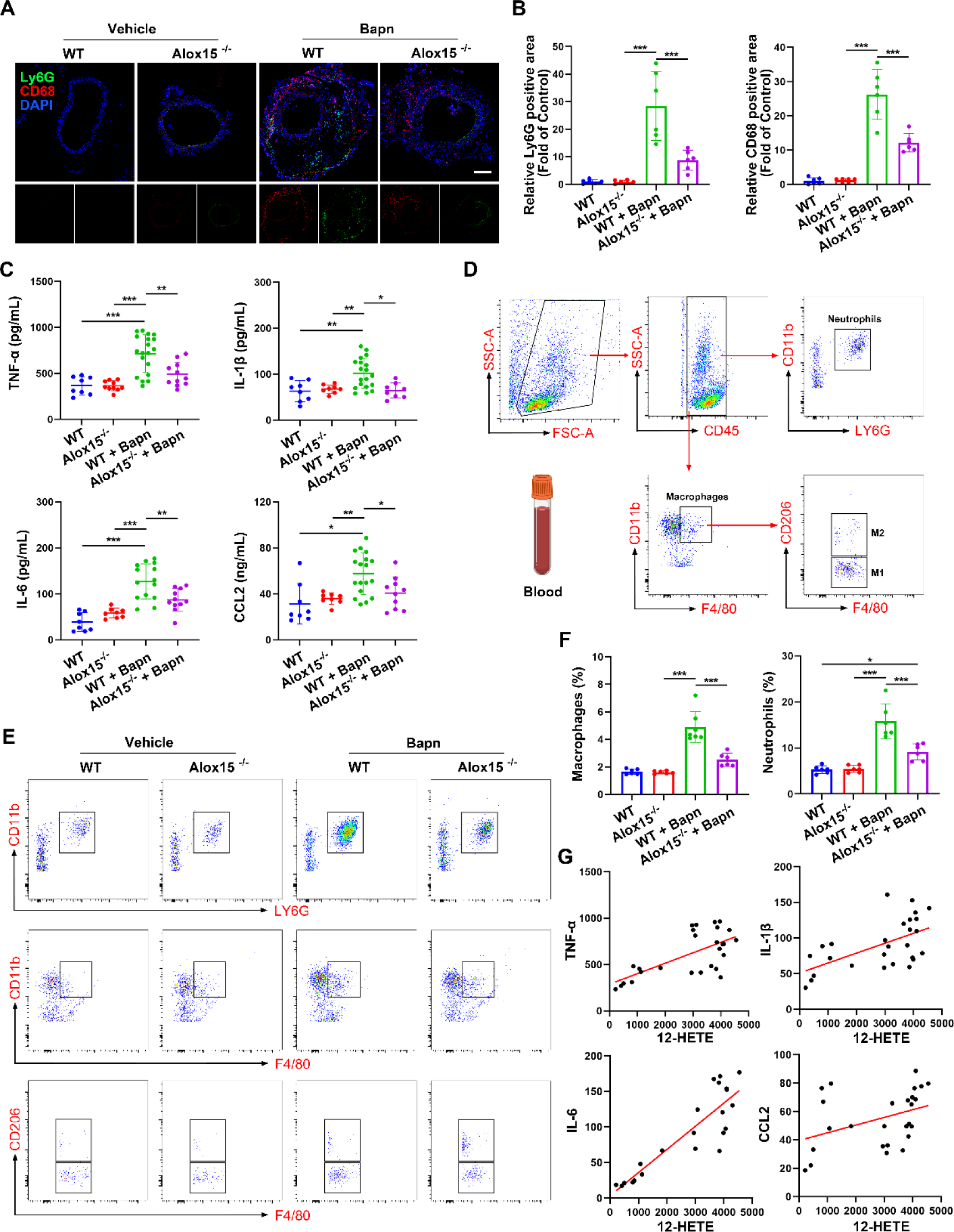
Alox15 deficiency reduces BAPN-induced vascular inflammation in mice. **A**, Representative images of frozen aorta sections stained with an antibody against Ly6G (green) and CD68 (red) nuclei were stained with DAPI (blue). Scale bar 50 μm. **B**, The statistical graph of the results is shown on the right (n = 6 in each group). The Ly6G-positive and CD68-positive areas were measured and expressed as the percentage of the positive area in sections out of the entire visual field of the section. **C**, ELISA analysis of inflammatory factors TNF-α, IL-1β, IL-6 and CCL2 in murine plasma at the endpoint (n = 6 each group). **D**, Gating strategy for identification of macrophages and neutrophils in mouse blood. macrophages were identified as CD45+CD11b+ F4/80+ and further classified as CD206+ and CD206-. neutrophils as CD45+CD11B+LY6G+. **E**, Representative flow cytometry analysis of circulating blood Ly6G+ CD11b+ neutrophils and CD11b + F4/80+ macrophages in BAPN - and saline-treated WT and Alox15^−/−^ mice. **F**, Flow cytometry Data analysis was performed as shown (n = 6 each group) G, Pearson correlation analysis of the serum 12HETE and serum inflammatory factors (TNF, IL1, IL6 and CCL2) of BAPN - and saline-treated WT and Alox15^−/−^ mice. \**p* < 0.05, \*\**p* < 0.01 and \*\*\**p* < 0.001, n.s. = not significant, one-way ANOVA, Tukey’s multiple comparisons test.

### 12-HETE is an Essential Driver of TAD Development Through the BLT2 Receptor in Mice

Because arachidonic acid can be converted to 12-HETE through a process mediated by the metabolic enzyme 12/15-LOX, it has been unclear whether 12-HETE participates in aortic dissection. We initiated intraperitoneal injection of 12-HETE in week 2 during BAPN-induced mouse modeling of TAD. As expected, 12-HETE injection exacerbated mortality in mice (n=19 and 23, Figure 5B). Additionally, the rates of aortic dissection formation and rupture increased from 95% (n =18) and 63% (n = 12) to 100% (n = 23) and 86% (n =20), respectively (Figure 5A and 5C). Vascular ultrasound images and measurements of maximum aortic diameter on day 28 after modeling also demonstrated that 12-HETE exacerbated BAPN-induced aortic dilation, compared with WT controls (Figure 5D and 5F). Hematoxylin and eosin staining and Elastica van Gieson staining showed that dissecting aneurysm formation and elastin disruption were worse in 12-HETE-treated mice than in saline-treated mice (Figure 5E and G). These findings indicated that the metabolite 12-HETE augmented TAD formation.

**Figure 5.**
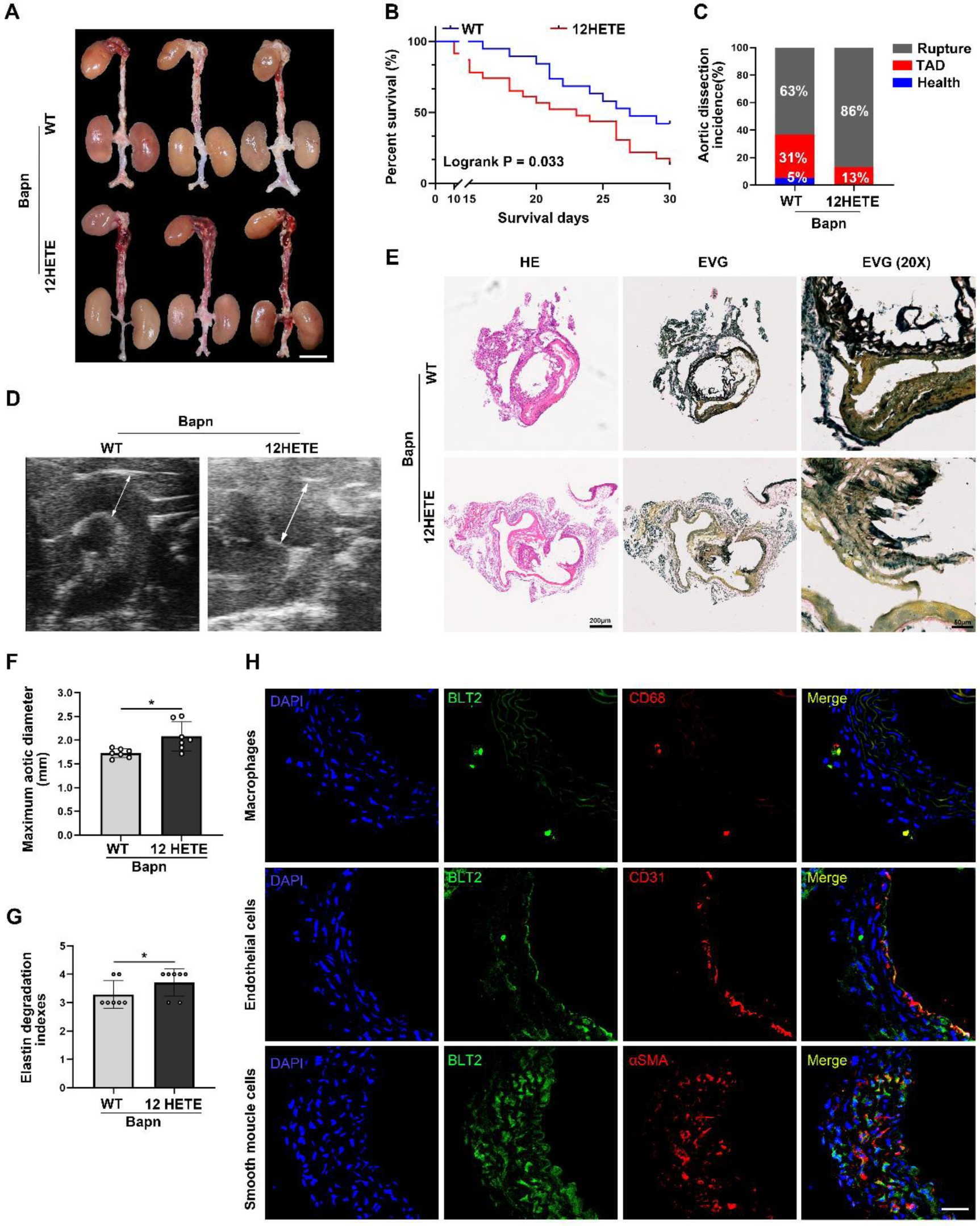
12 HETE is an essential driver of TAD development via BLT2 receptor in mice. **A** through **G**, C57BL/6 mice were observed with or without 12HETE after BAPN (β-aminopropionitrile monofumarate) treatment for 28 days. **A**, Representative macrographs of aorta (scale bar, 1 mm). **B**, Survival rate was estimated by Kaplan-Meier method and compared by log-rank test (P=0.0087; n=20 per group). **C**, Thoracic aortic dissection (TAD) incidence. **D**, Representative ultrasound images of thoracic aorta. **E**, Representative macroscopic images of aorta sections stained with hematoxylin and eosin (HE) and elastic– Van Gieson (EVG; scale bars, 100 μm). **F**, Measurements of maximum aortic diameter (n=9 per group). **G**, Elastin break grades were analyzed by Kruskal-Wallis followed by Dunn multiple comparisons test. **H**, Representative confocal images of BLT2 (green) colocalized with CD68 (red)–positive macrophages, CD31 (red) positive endothelial cells and αSMA (red) positive smooth muscle cells in aorta tissues from TAD mice. (scale bars, 50 μm). \**p* < 0.05, \*\**p* < 0.01 and \*\*\**p* < 0.001, n.s. = not significant, one-way ANOVA, Tukey’s multiple comparisons test.

BLT2, a receptor for 12-HETE, is reportedly expressed in endothelial cells (ECs). Here, we found that Alox15 was colocalized with the EC marker CD31, as well as the smooth muscle cell marker α-SMA and the macrophage marker CD68, in the aorta of TAD mice (Figure 5H). Mouse aortic endothelial cells (MAECs), primary smooth muscle cells, and primary macrophages also expressed BLT2 upon stimulation with angiotensin II (Ang II; Figure 6A and 6B). Furthermore, 12-HETE promoted BLT2 expression in primary SMCs and macrophages upon stimulation with Ang II (Figure 6D, 6E, and 6H). Single-cell sequencing of the aorta in healthy controls and TAD patients revealed that BLT2 mRNA was expressed in ECs, macrophages, SMCs, and T cells (Figure 6C, Figure S8).

**Figure 6.**
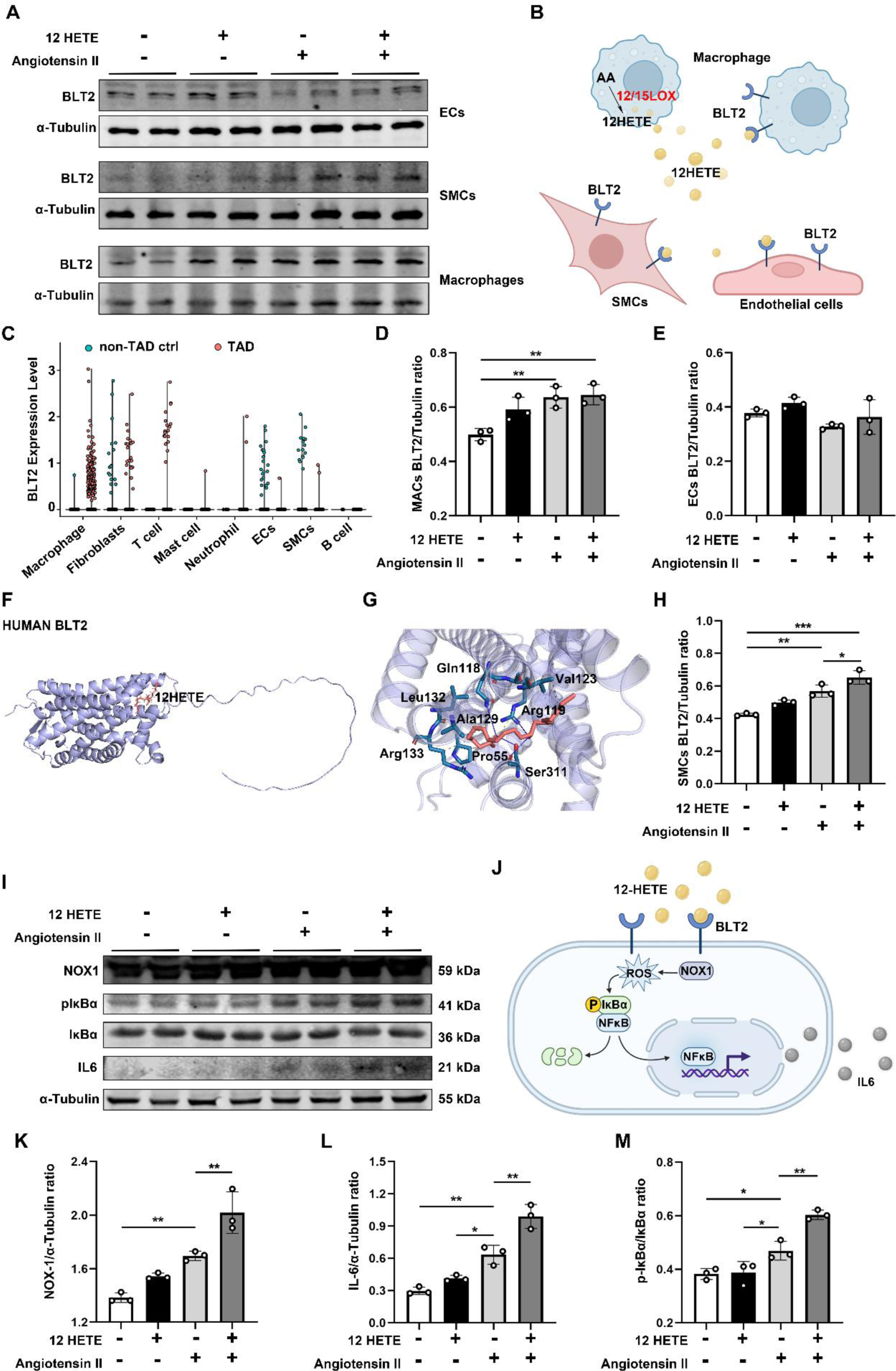
12 HETE Regulate VSMC phenotypic switch requires IL6 production. **A**, Representative Western blot images of BLT2 in MAECs, primary smooth muscle cells and primary macrophages. **B**, Graphical abstract. 12HETE combined with BLT2 expressed in macrophages, ECs and VSMCs. **C**, BLT2 mRNA expression from single-cell sequencing of human aorta. **D**, **E** and **H**, Western blot analysis and quantification of BLT2 expressed in macrophages, ECs and VSMCs. **F** and **G**, Representative docking images of 12HETE and BLT2 from human and mouse. **I**, Representative Western blot images of NOX-1, p-IκBb, IκBb and IL6 in primary macrophages. **J**, Graphical abstract. 12HETE induced NOX-1/ROS/NFκb activation via BLT2 receptor. **K**, **L** and **M**, Western blot analysis and quantification of NOX-1, p-IκBb/IκBb and IL6 expressed in macrophages. \**p* < 0.05, \*\**p* < 0.01 and \*\*\**p* < 0.001, n.s. = not significant, one-way ANOVA, Tukey’s multiple comparisons test.

Although BLT2 is a known receptor for 12-HETE, the interaction between 12-HETE and BLT2 has not been elucidated. Considering the importance of 12-HETE and BLT2 in TAD, we performed molecular docking analysis of 12-HETE and BLT2. This analysis showed that 12-HETE had robust binding affinity. 12-HETE forms intermolecular hydrogen bonds with amino acid residues of GLN, ARG, SER and SER and has hydrophobic interactions with PRO, VAL, VAL, ALA, LEU and ARG in BLT2 (Figure 6F and 6G). Taken together, these data suggest that 12-HETE is an essential driver of TAD development through the BLT2 receptor in mice.

### 12-HETE-Mediated Regulation of VSMC Phenotype Switching Requires IL-6 Production

Because 12-HETE and its receptor play critical roles in VSMC phenotype differentiation and inflammation onset, we explored the underlying molecular mechanism. It is unclear how the 12-HETE–BLT2 interaction affects VSMC differentiation. We speculated that 12-HETE binding to BLT2 activates downstream signaling. There is evidence that LTB4 receptors regulate ROS and inflammatory factors. Primary macrophages were stimulated with 12-HETE and/or Ang II. We observed that NOX-1 expression was promoted upon simulation with Ang II. Upon addition of 12-HETE, NOX-1 expression increased (Figure 6I–6K). 12-HETE and Ang II also enhanced the downstream conversion of p-IκBb to IκBb (Figure 6M). Finally, IL-6 was produced in large quantities upon activation of the NOX-1/ROS/NFκB signaling pathway (Figure 6L).

Furthermore, we stimulated mouse primary smooth muscle cells and VSMCs with various concentrations of IL-6. IF staining showed that increases in IL-6 concentrations led to reduced expression of α-SMA in both types of smooth muscle cells (Figure 7A–7C). We suspected that activation of the JAK/STAT signaling pathway was associated with phenotype switching in smooth muscle cells; thus, we performed WB to determine the expression levels of relevant proteins. The results showed that the levels of p-JAK/JAK and p-STAT/STAT were substantially upregulated with 10 ng/mL of IL-6 (Figure 7D–7H). In contrast, the expression levels of α-SMA and SM22α, proteins associated with phenotype switching in smooth muscle cells, were significantly downregulated (Figure 7I and 7J).

**Figure 7.**
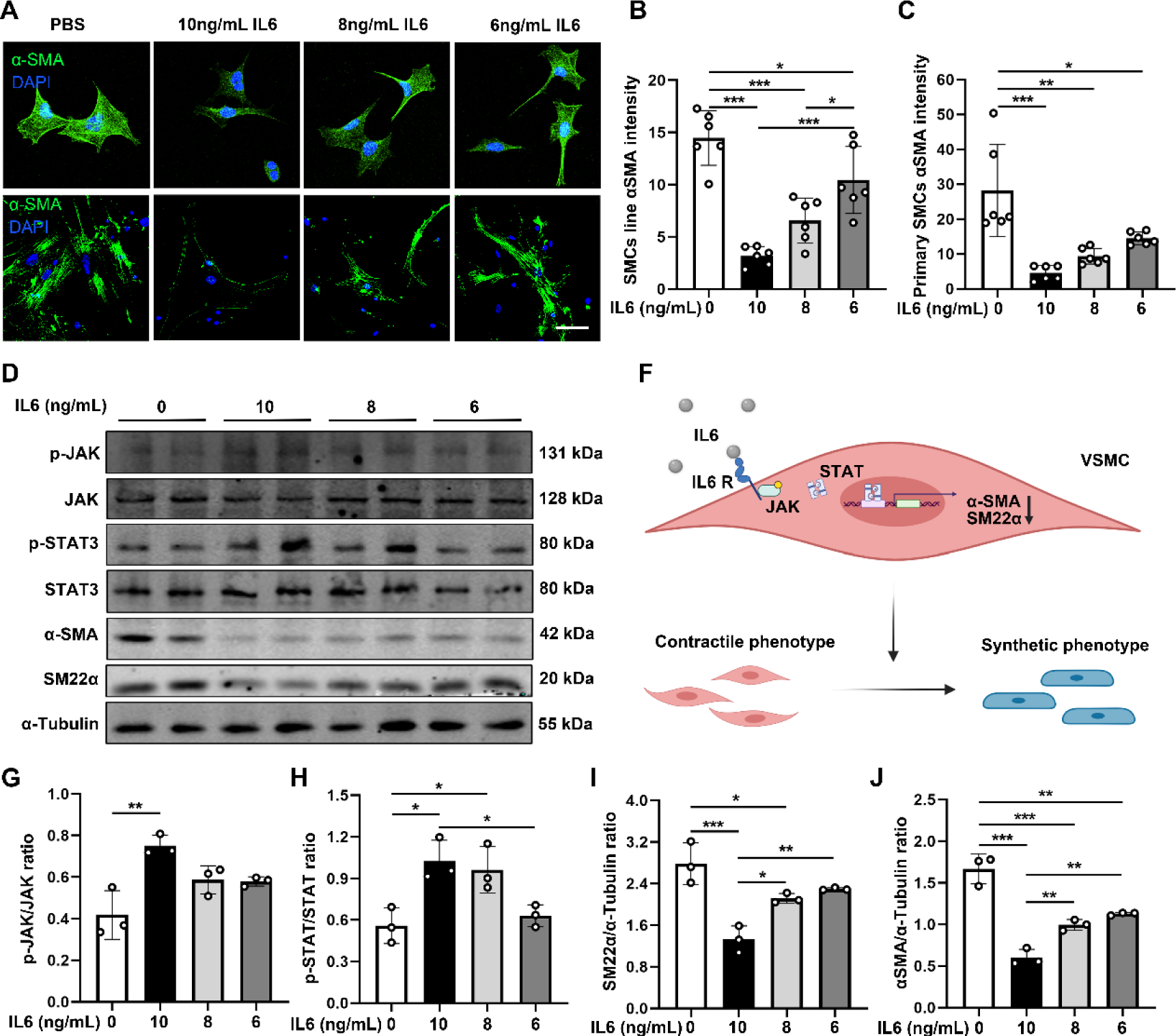
IL6 induced VSMC phenotypic switch though JAK/STAT signal pathway. **A**, Representative confocal images of αSMA (red) in primary smooth muscle cells and vascular smooth muscle cells. (scale bar, 50 μm). **B** and **C**, Quantification of αSMA levels from confocal images. **D**, Representative Western blot images of p-JAK, JAK, p-STAT, STAT, α-SMA and SM22α in primary macrophages. **F**, Graphical abstract. IL6 induced VSMC phenotypic switch though JAK/STAT. **G** – **J,** Western blot analysis and quantification of p-JAK/JAK, p-STAT/STAT, α-SMA and SM22α expressed in VSMCs. \**p* < 0.05, \*\**p* < 0.01 and \*\*\**p* < 0.001, n.s. = not significant, one-way ANOVA, Tukey’s multiple comparisons test.

### Pharmacologic Blockade of 12/15-LOX or Antagonism of the BLT2 Receptor Protects Mice from BAPN-Induced TAD

Through modulation of the Alox15/12-HETE/BLT2/NF-κB signaling pathway axis, we explored potential new targets for the clinical treatment of TAD. ML351, an inhibitor of 12/15-LOX, can effectively reduce 12/15-LOX expression, decreasing the serum level of 12-HETE. Additionally, LY255283 is a BLT2 receptor antagonist that competitively binds to elevated levels of 12-HETE in serum; this binding interaction hinders activation of the downstream NOX-1/ROS/NFκB/IL-6 signaling pathway, thereby alleviating the inflammatory response associated with TAD. We explored the therapeutic effects of ML351 and LY255283 on TAD development in a mouse model of BAPN-induced TAD. C57BL/6J mice were intraperitoneally injected with ML351 and/or LY255283 daily after BAPN treatment throughout the 4-week modeling period. As anticipated, the ML351 and LY255283 groups (n=22) showed reduced TAD formation (Figure 8A and 8D) and lethality (Figure 8C and 8D) as well as attenuated aortic dilation (Figure 8D and 8E), compared with the control group. When ML351 and LY255283 were used in combination, the rates of TAD incidence and rupture were greatly reduced, and the mortality rate was decreased. TAD progression was significantly inhibited. Histological detection by hematoxylin and eosin staining and Elastica van Gieson staining revealed identical pathological changes, in accordance with previous findings that 12/15-LOX inhibition and BLT2 antagonism rescued elastin disorganization and vessel wall dissection (Figure 8F). These results revealed the central roles of 12/15LOX, 12-HETE and BLT2 in TAD progression, indicating that ML351 and LY255283 are promising drugs for the treatment of TAD.

**Figure 8.**
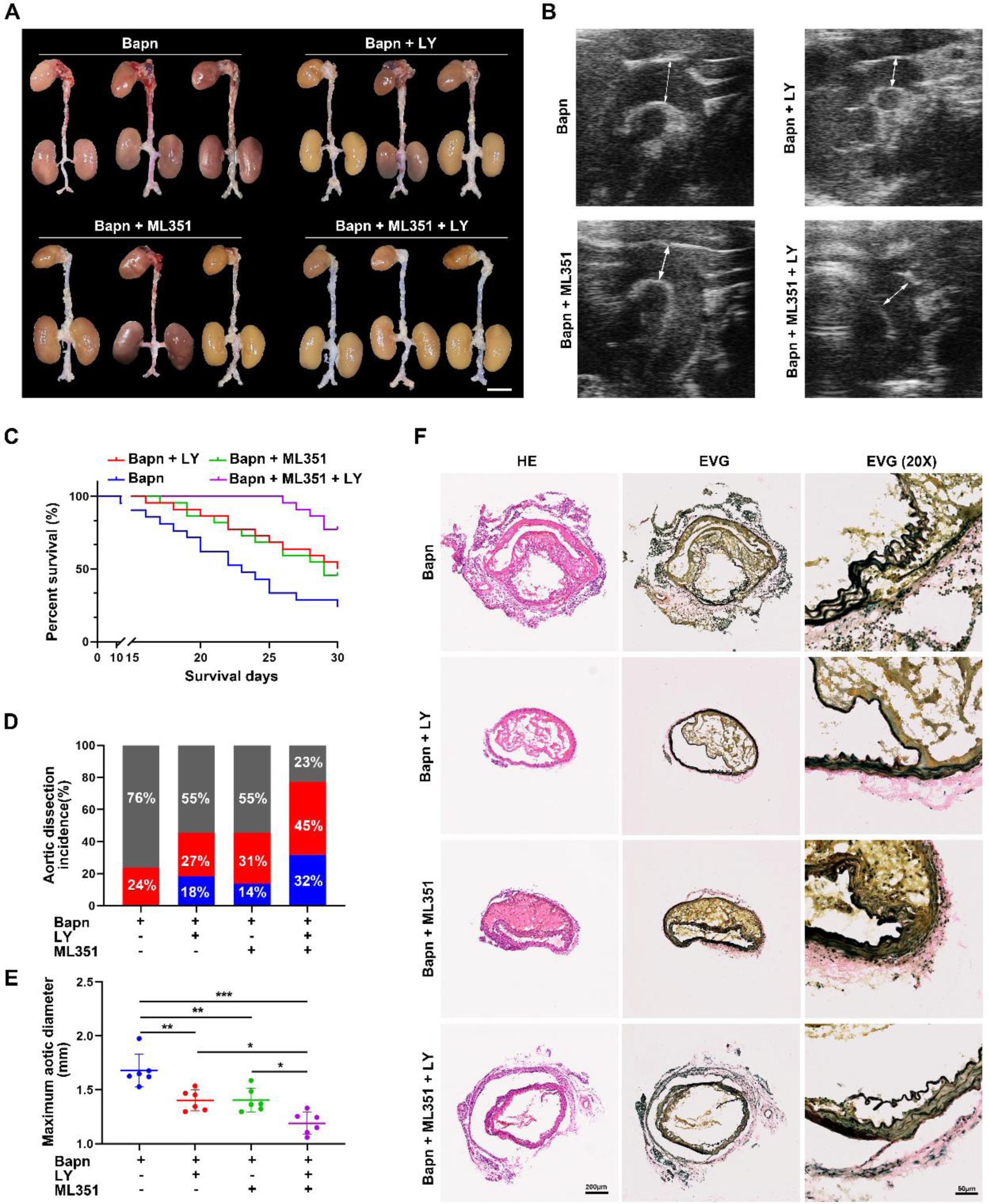
Pharmacologic blockade of Alox15 or antagonizing BLT2 receptor Protects mice from BAPN-induced TAD. **A** through **G**, C57BL/6 mice were observed with or without ML351 and LY255283 after BAPN (β-aminopropionitrile monofumarate) treatment for 28 days. **A**, Representative macrographs of aorta (scale bar, 1 mm). **B**, Survival rate was estimated by Kaplan-Meier method and compared by log-rank test (n=20 per group). **C**, Thoracic aortic dissection (TAD) incidence. **D**, Representative ultrasound images of thoracic aorta. **E**, Representative macroscopic images of aorta sections stained with hematoxylin and eosin (HE) and elastic– Van Gieson (EVG; scale bars, 200 μm; 50 μm). **F**, Measurements of maximum aortic diameter (n=9 per group). **G**, Elastin break grades were analyzed by Kruskal-Wallis followed by Dunn multiple comparisons test. \**p* < 0.05, \*\**p* < 0.01 and \*\*\**p* < 0.001, n.s. = not significant, one-way ANOVA, Tukey’s multiple comparisons test.

## Discussion

In this study, we observed increased levels of 12-HETE in serum as well as elevated 12/15-LOX expression in diseased aortic tissue from TAD patients and TAD mice. Genetic and pharmacologic blockade of Alox15 reduced 12-HETE released, suppressed inflammation, and VSMC phenotype switching in TAD mice. Moreover, we confirmed that 12-HETE triggered an inflammatory cascade and facilitated VSMC phenotype switching by binding to the BLT2 receptor in macrophages and VSMCs. Thus, blockade of BLT2 prevented activation of the NOX-1/ROS/NFκB/IL-6 signaling pathway, downregulated VSMC differentiation markers through the JAK/STAT signaling pathway, and eventually attenuated TAD development. Our results indicate that 12-HETE is a trigger for inflammatory cascade, a mechanism highly relevant to aberrant VSMC phenotype switching. Inhibition of 12-HETE-related pathways (e.g., lowering plasma 12-HETE levels or blocking its receptor) decreased TAD risk.

Arachidonic acid, an integral part of all cell membranes is released from membrane phospholipids following mechanical stimulation or stress.^17, 18^ Arachidonic acid can be metabolized into numerous metabolites via cyclooxygenase (COX), lipoxygenase (LOX), and cytochrome P450 (CYP450) with different functions and activities.^19^ By using targeted metabolomics analysis of eicosanoids, we identified that the active metabolites of the LOX pathway were elevated in TAD patients and mice including 12-HETE, 8-HETE, and 15-HETE. Among them, 12-HETE showed the most significant fold change. Therefore, we sought to determine the origin of 12-HETE production in TAD patients and mice. Analysis of diseased aortic tissue revealed elevated levels of 12/15-LOX in TAD patients and TAD mice. Accordingly, IF staining demonstrated increased 12/15-LOX colocalization in macrophages in cardiac tissue. Consistent with findings of previous single-cell RNA-Seq data of aortic dissection tissue harvest from BAPN-induced TAD mice, they demonstrated that Alox15 increased in a subpopulation of macrophage as well as SMC.^20^ Therefore, we speculate that macrophage-derived 12/15-LOX is a source of the elevated serum 12-HETE levels in TAD, observed in response to early vascular injury though more experiments need for further verification.

To further explore how 12-HETE, the major metabolite of leukocyte-specific 12/15-LOX, was involved in the development of TAD, we constructed an Alox15 knockout mouse model and determined that deletion of 12/15LOX decreased serum 12-HETE, improved survival and reduced rupture rates. Meanwhile, 12-HETE supplementation may aggravate aortic dissection in mice. Inappropriate enhanced function of Alox15 and 12-HETE is known to trigger an inflammatory response that exacerbates organ damage.^21^ We found that 12-HETE increased before inflammatory cytokines at an early stage of TAD development after 3 weeks of BAPN administration. Besides, 12-HETE levels were positively related to inflammatory cytokines including IL-6 in TAD, the latter rapidly activates JAK/STAT in vascular smooth muscle cells to promote an inflammatory synthetic phenotype. Therefore, 12-HETE is paramount to the initiation and propagation of inflammation and subsequent VSMC phenotype switching in TAD development. 12-HETE has been discovered as the endogenous ligand for BLT2.^22, 23^ Our findings confirmed the colocalization of 12-HETE with BLT2 in macrophages, ECs, and VSMCs. The binding of 12-HETE to BLT2 results in the activation of BLT2, thereby stimulating the activation of the NOX-1/ROS/NFκB/IL-6 signaling pathway in macrophages, ECs, and VSMCs. The BLT2 inhibitor LY255283 reversed the protective effect of Alox15 deletion on TAD progression. BLT2 is a known member of the LTB4 family.^24, 25^ However, without avaliable structure of BLT2, their specific binding sites and mechanism of action remain unclear. We use Alpha Fold model to predict the protein structure of BLT2 and examinate the binding affinity and mode of binding between 12-HETE and BLT2 using molecular docking analysis. Collectively, we identified putative binding sites for 12-HETE and BLT2 and provides evidence that 12-HETE activated the downstream NOX-1/ROS/NFκB/IL-6 signaling pathway, leading to inflammatory factor production and promoting VSMC phenotype switching via BLT2 activation.

Our study identified that 12-HETE initiates the inflammatory cascade and triggers aberrant phenotype switching in VSMCs via BLT2 receptor during TAD development and shows potential as a biomarker of TAD. However, further validation of its clinical utility in TAD risk prediction is still required in following clinical trials in cohorts. Besides, to our knowledge, at this point, it is still unclear which BLT2 domain binds 12-HETE. Although molecular docking experiments were performed to predict the binding site of 12-HETE to a predicted BLT2 protein structure by Alphafold, our result needs further confirmation in cryo-electron microscopy ultrastructural studies. The combination of the 12/15-LOX inhibitor and BLT2 blocker greatly reduces aortic injury in TAD mice. But the chronic safety and systematic influence of both inhibitors should be investigated before considering the clinical success of TAD prevention by pharmaceutically targeting this target.

The present study demonstrated the effects of 12/15-LOX and its metabolite 12-HETE on TAD development. During TAD progression, 12-HETE triggers the initiation of an inflammatory cascade and substantial VSMC differentiation through binding to BLT2 on macrophages. Targeted inhibition of 12/15-LOX and blockade of BLT2 may be useful in the development of novel therapeutic strategies.

## Acknowledgments

We thank Ryan Chastain-Gross, Ph.D., from Liwen Bianji (Edanz) (www.liwenbianji.cn/) for editing the English text of a draft of this manuscript.

## Authors’ contributions

The contribution of each author is substantiated as followed: Conceived and designed the research: Yuan Wang, and Jie Du; Acquired the data: Yuyu Li, Jiaqi Yu, Weiyao, Rui Lin, Xue Wang and Wenxi Jiang; Performed mass spectrometry analysis: Xin Tan and Xuan Xu, Performed statistical analysis: Yuyu Li, Jiaqi Yu and Weiyao Chen; Drafted the manuscript: Yuyu Li, Jiaqi Yu and Weiyao Chen; Made critical revision of the manuscript for key intellectual content: Yuan Wang and Jie Du.

## Reference

1. Rylski, B.; Schilling, O.; Czerny, M. European heart journal 2023, 44, (10), 813–821.

2. Goldfinger, J. Z.; Halperin, J. L.; Marin, M. L.; Stewart, A. S.; Eagle, K. A.; Fuster, V. Journal of the American College of Cardiology 2014, 64, (16), 1725–39.

3. Bortone, A. S.; De Cillis, E.; D’Agostino, D.; de Luca Tupputi Schinosa, L. Circulation 2004, 110, (11 Suppl 1), Ii262–7.

4. Chou, E.; Pirruccello, J. P.; Ellinor, P. T.; Lindsay, M. E. Nature reviews. Cardiology 2023, 20, (3), 168–180.

5. El-Hamamsy, I.; Yacoub, M. H. Nature reviews. Cardiology 2009, 6, (12), 771–86.

6. Chakraborty, A.; Li, Y.; Zhang, C.; Li, Y.; Rebello, K. R.; Li, S.; Xu, S.; Vasquez, H. G.; Zhang, L.; Luo, W.; Wang, G.; Chen, K.; Coselli, J. S.; LeMaire, S. A.; Shen, Y. H. Circulation 2023, 148, (12), 959–977.

7. Branchetti, E.; Poggio, P.; Sainger, R.; Shang, E.; Grau, J. B.; Jackson, B. M.; Lai, E. K.; Parmacek, M. S.; Gorman, R. C.; Gorman, J. H.; Bavaria, J. E.; Ferrari, G. Cardiovascular research 2013, 100, (2), 316–24.

8. Meng, H.; Liu, Y.; Lai, L. Accounts of chemical research 2015, 48, (8), 2242–50.

9. Bensinger, S. J.; Tontonoz, P. Nature 2008, 454, (7203), 470–7.

10. Davies, P.; Bailey, P. J.; Goldenberg, M. M.; Ford-Hutchinson, A. W. Annual review of immunology 1984, 2, 335–57.

11. Ma, X. H.; Liu, J. H.; Liu, C. Y.; Sun, W. Y.; Duan, W. J.; Wang, G.; Kurihara, H.; He, R. R.; Li, Y. F.; Chen, Y.; Shang, H. Signal transduction and targeted therapy 2022, 7, (1), 288.

12. Cai, W.; Liu, L.; Shi, X.; Liu, Y.; Wang, J.; Fang, X.; Chen, Z.; Ai, D.; Zhu, Y.; Zhang, X. Circulation 2023, 147, (19), 1444–1460.

13. He, L.; Lin, Y.; Wang, X.; Liu, X. L.; Wang, Y.; Qin, J.; Wang, X.; Day, D.; Xiang, J.; Mo, J.; Zhang, Y.; Zhang, J. J. Environment international 2020, 145, 106154.

14. Fang, K. M.; Lee, A. S.; Su, M. J.; Lin, C. L.; Chien, C. L.; Wu, M. L. Cardiovascular research 2008, 78, (3), 533–45.

15. Ringnér, M. Nature biotechnology 2008, 26, (3), 303–4.

16. Pernet, E.; Sun, S.; Sarden, N.; Gona, S.; Nguyen, A.; Khan, N.; Mawhinney, M.; Tran, K. A.; Chronopoulos, J.; Amberkar, D.; Sadeghi, M.; Grant, A.; Wali, S.; Prevel, R.; Ding, J.; Martin, J. G.; Thanabalasuriar, A.; Yipp, B. G.; Barreiro, L. B.; Divangahi, M. Nature 2023, 614, (7948), 530–538.

17. Samuelsson, B.; Dahlén, S. E.; Lindgren, J. A.; Rouzer, C. A.; Serhan, C. N. Science (New York, N.Y.) 1987, 237, (4819), 1171–6.

18. Bazinet, R. P.; Layé, S. Nature reviews. Neuroscience 2014, 15, (12), 771–85.

19. Tigistu-Sahle, F.; Lampinen, M.; Kilpinen, L.; Holopainen, M.; Lehenkari, P.; Laitinen, S.; Käkelä, R. Journal of lipid research 2017, 58, (1), 92–110.

20. Liu, X.; Chen, W.; Zhu, G.; Yang, H.; Li, W.; Luo, M.; Shu, C.; Zhou, Z. Cell discovery 2022, 8, (1), 11.

21. Lu, C. L.; Teng, T. Y.; Liao, M. T.; Ma, M. C. International journal of molecular sciences 2021, 22, (12).

22. Ro, M.; Lee, A. J.; Kim, J. H. Allergy 2018, 73, (2), 350–360.

23. Kriska, T.; Herrnreiter, A.; Pfister, S. L.; Adebesin, A.; Falck, J. R.; Campbell, W. B. Hypertension (Dallas, Tex. : 1979) 2022, 79, (1), 104–114.

24. Brink, C.; Dahlén, S. E.; Drazen, J.; Evans, J. F.; Hay, D. W.; Nicosia, S.; Serhan, C. N.; Shimizu, T.; Yokomizo, T. Pharmacological reviews 2003, 55, (1), 195–227.

25. Yokomizo, T.; Kato, K.; Terawaki, K.; Izumi, T.; Shimizu, T. The Journal of experimental medicine 2000, 192, (3), 421–32.

